# CD18-mediated adhesion is required for lung inflammation induced by mononuclear cell-derived extracellular vesicles

**DOI:** 10.1101/158121

**Authors:** Tommaso Neri, Valentina Scalise, Ilaria Passalacqua, Ilaria Giusti, Cristina Balia, Delfo D’Alessandro, Stefano Berrettini, Roberto Pedrinelli, Pierluigi Paggiaro, Vincenza Dolo, Alessandro Celi

**Affiliations:** Laboratorio di Biologia Cellulare Respiratoria; Dipartimento di Patologia Chirurgica, Medica, Molecolare e dell’Area Critica. University of Pisa, and Azienda Ospedaliero-Universitaria Pisana, Pisa, Italy; Department of Life, Health and Environmental Sciences, University of L’Aquila, L’Aquila, Italy; OtoLab; Dipartimento di Patologia Chirurgica, Medica, Molecolare e dell’Area Critica. University of Pisa, and Azienda Ospedaliero-Universitaria Pisana, Pisa, Italy

**Keywords:** Extracellular vesicles, integrins, lung inflammation, adhesion, CD-18

## Abstract

Extracellular vesicles are submicron vesicles that upregulate the synthesis of proinflammatory mediators by lung epithelial cells.

We investigated whether these structures adhere to lung epithelial cells, and whether adhesion is a prerequisite for their proinflammatory activity.

Extracellular vesicles were generated by stimulation of normal human mononuclear cells with the calcium ionophore A23187, and labelled with carboxyfluorescein diacetate succinimidyl ester. Adhesion of vesicles to monolayers of immortalized bronchial epithelial cells (16HBE) and alveolar cells (A549) was analysed by fluorescence microscopy. The role of candidate adhesion receptors was evaluated with inhibitory monoclonal antibodies and soluble peptides. The synthesis of proinflammatory mediators was assessed by ELISA.

Transmission electron microscopy confirmed the generation of closed vesicles with an approximate size range between 50 and 600 nm. Adhesion of extracellular vesicles to epithelial cells was minimal in baseline conditions and was upregulated upon stimulation of the latter with tumour necrosis factor-α. Adhesion was blocked by an anti-CD18 antibody and by peptides containing the sequence RGD. The same molecules also blocked the upregulation of the synthesis of interleukin-8 and monocyte chemotactic protein-1 induced by extracellular vesicles.

**Summary statement:** Extracellular vesicles upregulate the synthesis of proinflammatory mediators by lung epithelial cells. CD18-mediated adhesion to target cells is required for this proinflammatory effect and might represent a target for anti-inflammatory therapy.

## Introduction

Extracellular vesicles (EV) are submicron structures shed by cells under various conditions (van der Pol et al., 2012). A growing body of evidence indicates that these previously underappreciated structures are involved in numerous pathophysiologically relevant phenomena including, for example, blood coagulation and thrombosis (Celi et al., 2004; Geddings and Mackman, 2014), inflammation (Andriantsitohaina et al., 2012), cancer development and progression (Giusti and Dolo, 2014; Żmigrodzka et al., 2016), neurological disorders (Yun et al., 2016). The role of EV in pulmonary diseases is also gaining consideration (Nieri et al., 2016).

EV exert their biological effects through different mechanisms. The presence of phosphatidylserine on their outer membrane of some types of EV accounts, at least in part, for their procoagulant potential (Owens and Mackman, 2011). However, EV also carry on their surface, and contain within their cytoplasm, some of the molecules present in the parental cell, including integral membrane proteins, soluble mediators, nucleic acids, micro-RNAs (Pitt et al., 2016). The type and abundance of such molecules depend on the stimulus that caused EV generation (Jimenez et al., 2003; Bernimoulin et al., 2009).

We have shown that EV derived from human monocytes/macrophages stimulated with the calcium ionophore A23187 and with histamine upregulate lung epithelial cell synthesis of proinflammatory cytokines, namely interleukin (IL)-8 and monocyte chemotactic protein (MCP)-1, and of the leukocyte adhesion molecule intercellular adhesion molecule (ICAM) - 1 (Cerri et al., 2006). We have also shown that the effect is mediated by the activation of the nuclear factor (NF)-κB (Neri et al., 2011). Other groups have demonstrated that EV of different origin stimulate cytokine synthesis by endothelial cells as well (Wang et al., 2011), in some cases with NF-κB-independent mechanisms (Mesri and Altieri, 1999). All in all, the mechanisms of EV-mediated activation of target cells are complex and largely unknown. One key question that has not been fully addressed is whether EV must adhere to target cells in order to induce an inflammatory phenotype.

The aim of the present study was to investigate whether mononuclear cell-derived EV adhere to lung epithelial cells, and to test the hypothesis that EV adhesion is a prerequisite for their proinflammatory activity.

## Results

### Microscopic characterization of EV

In previous publications, we selected specific subtypes of EV (namely the so-called microparticles) by a combination of methods, including flow cytometry, NanoSight technology and analysis of PS content (Neri et al., 2016b; Petrini et al., 2016; Neri et al., 2016a). Because the data presented herein are obtained with material that could not be preemptively selected based on any of the indicated methods, we preliminarily analyzed it by TEM. As shown in figure 1 (panels A-C), the material used to stimulate epithelial cells consists of closed rounded vesicles that appear delimited by a white membrane on a dark background. Most vesicles range in size from 50 up to 180 nm indicating that the vesicles likely represent the structures commonly referred to as exosomes and microparticles (Nieri et al., 2016). A few larger vesicles (up to approximately 600 nm) were also observed (not shown).

**Figure 1.**
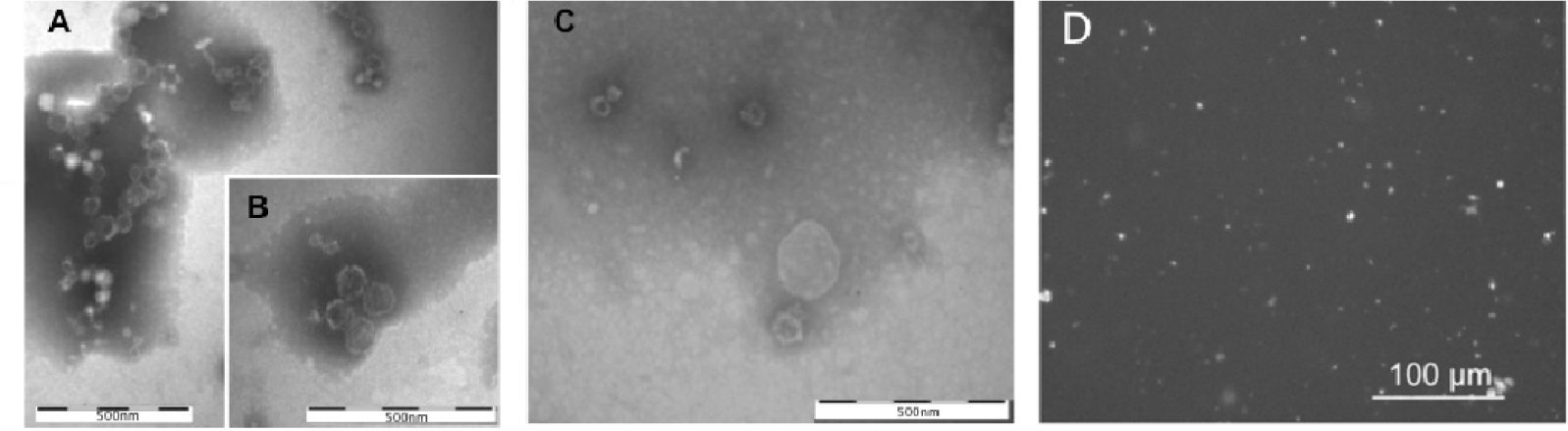
Microscopic characterization of EV. TEM micrographs of EV (A-C); bar: 500 nm. D. Fluorescence micrograph of CFSE-labelled EV prior to incubation with epithelial cells; bar: 100 μm.

### Mononuclear cell-derived EV adhere to lung epithelial cells

To investigate whether EV adhere to lung epithelial cells, EV generated by mononuclear cells stimulated with the calcium ionophore A23187 were labelled with CFSE and incubated at 37°C with A549 and 16HBE cells in the presence of Ca^++^ ions. Figure 1C shows the appearance of CFSE labelled EV prior to their incubation with epithelial cells. As most extracellular vesicles would be smaller than the resolution power of the optical microscope, the fluorescent spots likely represent clusters of vesicles as demonstrated by TEM. After a 90-min incubation, the number of EV adhering to unstimulated cells was low. Prestimulation of epithelial cells with TNF-α (25 ng/mL for 18 h) increased EV adhesion (fig. 2). The increase in EV adhesion after TNF-α stimulation of epithelial cells was statistically significant for both A549 and 16HBe cells (fig. 3A and 3B, respectively). Upon longer coincubations (18 hours), EV adhesion was significantly increased even in the absence of TNF-α prestimulation (fig. 3C and D).

**Figure 2.**
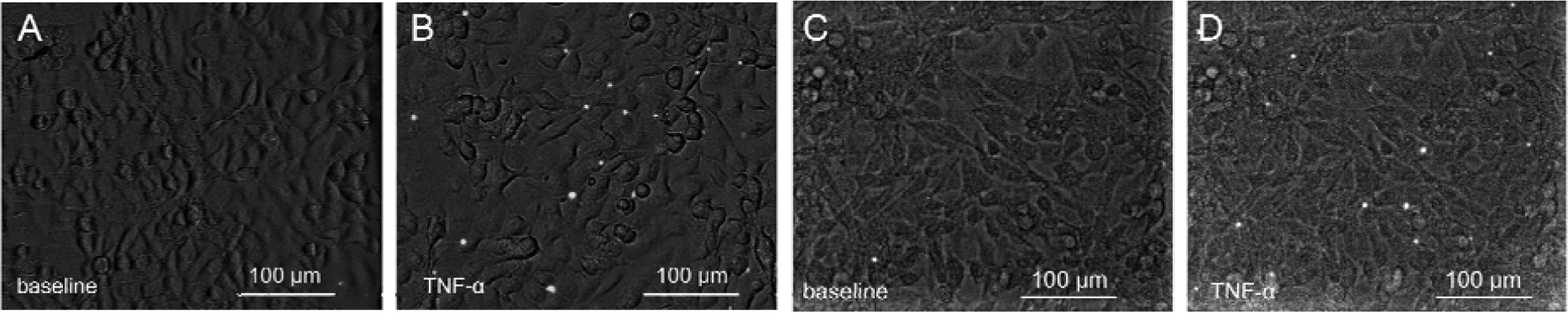
Adhesion of EV to lung epithelial cells. EV labelled with CFSE (A) were incubated with unstimulated (B) or TNF-α-stimulated (C) A549 cells and unstimulated (D) or TNF-α-stimulated (E) 16HBE cells for 1.5 hours at 37°C. After extensive washing, fluorescent events were enumerated. Images were obtained overlapping dark field and fluorescent field using ImageJ 1.49V and enhanced with Pixelmator 3.6 (Vilnius, Lithuania) using identical settings for panels B and C and for panels D and E. Bar: 100 μm. All images are representative of 10 consecutive experiments.

**Figure 3.**
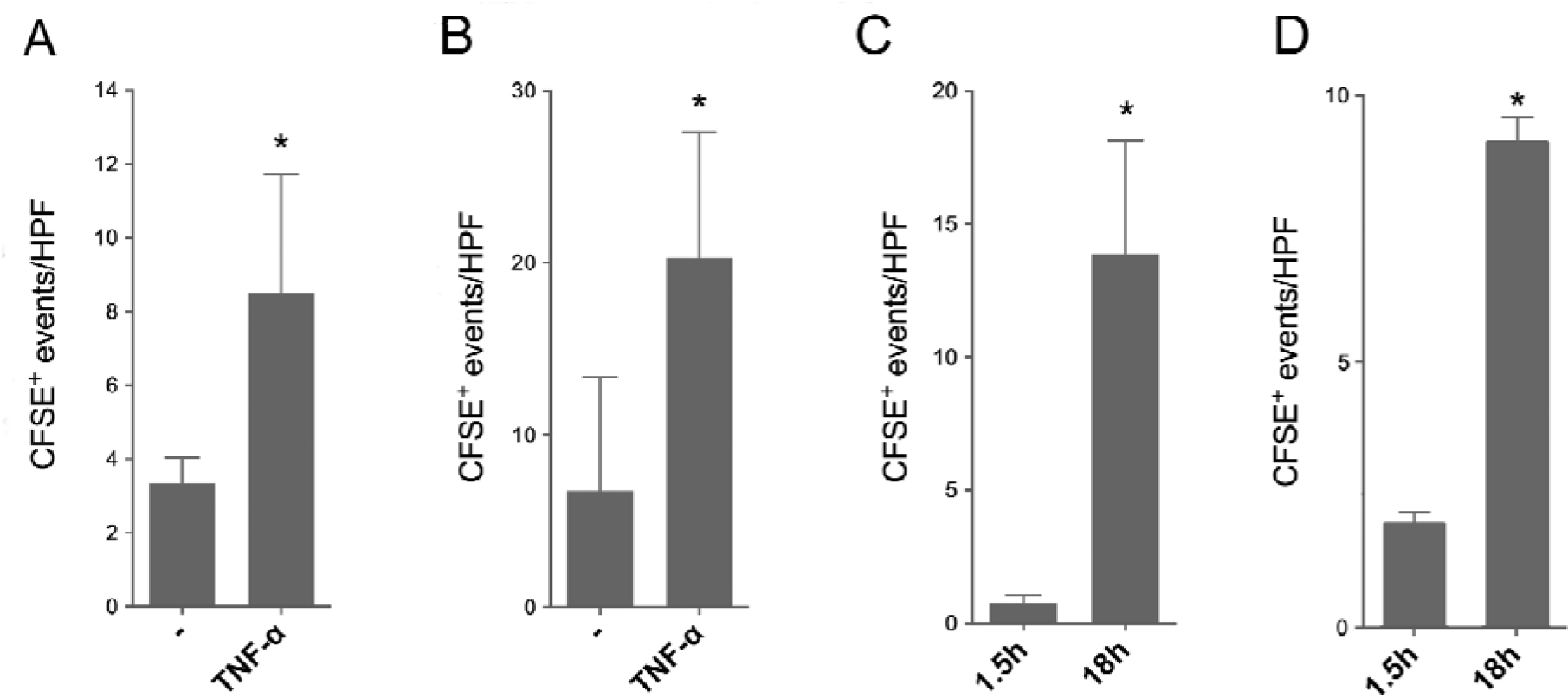
Adhesion of EV to lung epithelial cells in different conditions. EV were incubated with lung epithelial cells and enumerated as described in figure 2. A and B: Adhesion of EV to unstimulated and TNF-α-stimulated A549 and 16HBE cells, respectively, after 1.5-hour incubation. Data from 10 and 6 consecutive experiments for panel A and B, respectively; *: p<.05 for TNF-α-stimulated vs. unstimulated cells; paired t-test. C and D: Adhesion of EV to unstimulated A549 and 16HBE cells, respectively, after 1.5-and 18-hour incubations. *: p<.05 for 18-hour incubation vs. 1.5 hour incubation; paired t-test. Data from 4 and 3 consecutive experiments for panel C and D, respectively.

### Metal ions dependence of EV adhesion to TNFα stimulated epithelial cells

To begin to elucidate the mechanisms of EV adhesion, the experiments described in the previous paragraph were repeated under different conditions. First, we investigated the requirement for metal ions. As shown in figure 4A, either Ca^++^ or Mg^++^ were sufficient to fully support adhesion as the combination of the two ions did not further increase adhesion. In the presence of Mn^++^, EV adhesion was nearly maximal even in the absence of prior TNF-α stimulation (4B), mimicking results obtained with leukocytes (Dransfield et al., 1992). When experiments were carried out at different temperatures, we observed that adhesion was low at 4°C (fig. 4C).

**Figure 4.**
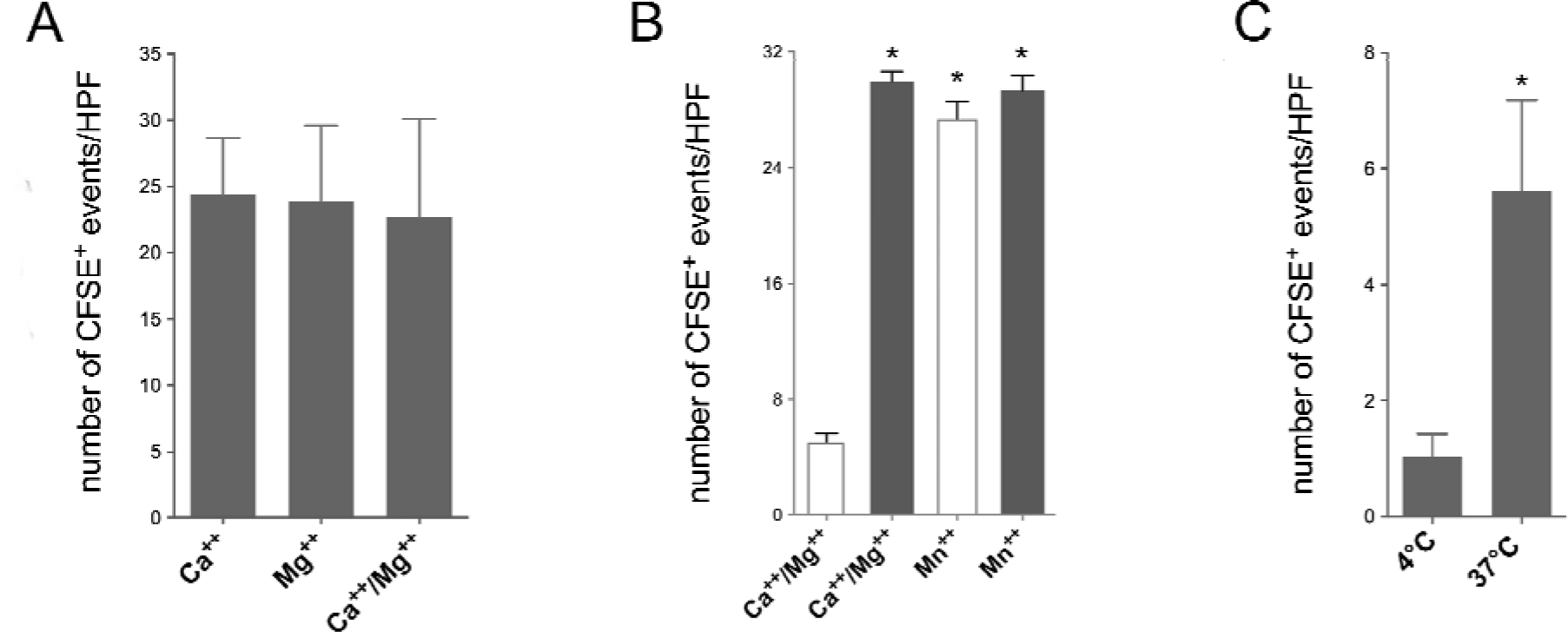
Adhesion of EV to lung epithelial cells in the presence of different metal ions and at different temperatures. EV were incubated with A549 cells for 1.5 hours and enumerated as described in figure 1. A. Adhesion of EV to TNF-α-stimulated A549 cells in the presence of Ca^++^ only, Mg^++^ only and Ca^++^ and Mg^++^. B. Adhesion of EV to unstimulated (open bars) or TNF-α-stimulated (solid bars) A549 cells in the presence of Ca^++^ and Mg^++^ or Mn^++^ - *: p<.05 for TNF-α-stimulated vs. unstimulated cells (in the presence of Ca^++^and Mg^++^) and for TNF-α-stimulated and unstimulated cells in the presence of Mn^++^ vs. unstimulated cells in the presence of Ca^++^and Mg^++^; repeated measures ANOVA. C. Adhesion of EV to TNF-α-stimulated A549 cells after incubation at 4°C and 37°C. *: p<.05 for adhesion at 37°C vs. 4°C; paired t-test. Data from 5, 5 and 4 consecutive experiments for panel A, B and C, respectively.

### Role of CD18 and RGD peptides on EV adhesion to TNF-α stimulated lung epithelial cells

We investigated the hypothesis that EV adhesion to lung epithelial cells is mediated by the integrin (β _2_subunit, CD18. Preincubation of EV with the inhibitory anti CD18 monoclonal antibody, TS1/18, inhibited adhesion in a dose dependent fashion (fig. 5A). Under our experimental conditions, the inhibitory effect of TS1/18 reached near maximum at 25 μg/mL. The non-immune, isotype-matched antibody, MOPC-21, did not cause any clear inhibition at concentrations up to 50 μg/mL (not shown). Similar results were obtained with 16HBE (fig. 5B). A peptide comprising the sequence arginine-glycine-aspartic acid (GRGDNP) dose dependently inhibited the adhesion of EV to A549 cells (fig. 5C). In contrast, the control peptide containing the scrambled sequence (GRADSP) had no effect (not shown).

**Figure 5.**
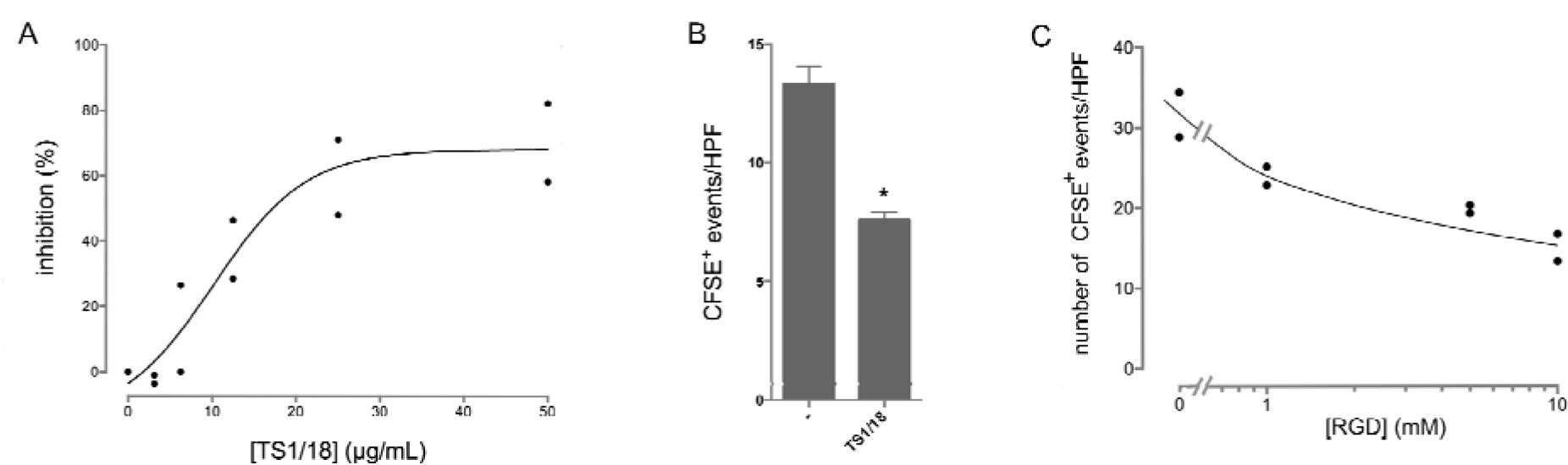
The anti-CD18 monoclonal antibody, TS1/18 and an RGD-containing peptide inhibit EV adhesion to lung epithelial cells. A. Dose-response inhibition of EV adhesion to TNF-α-stimulated A549 cells by TS1/18. B. Data from one experiment representative of 2. Inhibition of EV adhesion to TNF-α-stimulated 16HBE cells by TS1/18 (25 μg/mL). *: p<.05 for adhesion in the presence of the anti-CD18 antibody TS1/18 vs. adhesion in the absence of the antibody; paired t-test. Data from 4 consecutive experiments; C. Dose-response inhibition of EV adhesion to TNF-α-stimulated A549 cells by a peptide containing the sequence RGD. Data from one experiment representative of 4

### Adhesion to epithelial cells is required for EV-mediated upregulation of proinflammatory mediator synthesis

We have previously demonstrated that EV derived from mononuclear cells upregulate the synthesis of proinflammatory mediators, including IL-8 and MCP-1, by lung epithelial cells (Cerri et al., 2006). However, whether EV must adhere to epithelial cells for this effect to take place is not known. To address this issue, A549 and 16HBE cells were incubated with EV in the presence of the anti-CD18 antibody, TS1/18. As shown in figure 5, preincubation of EV with TS1/18 inhibited the expression of IL8 (6A) and MCP-1 (6B) by A549 cells; the effect was statistically significant. The dose-dependency of this effect parallels the inhibition of adhesion. MOPC-21 did not cause any clear inhibition (not shown). Similar results were obtained with 16HBE cells (fig. 6C). MCP-1 was not upregulated in 16HBE cells by mononuclear cell-derived EV (not shown) confirming previous data with normal human bronchial epithelial cells (Cordazzo et al., 2014). Figure 7 shows that both IL-8 and MCP-1 synthesis by A549 cells were also inhibited by an RGD-containing peptide. Again, the control scrambled peptide had no effect (not shown).

**Figure 6.**
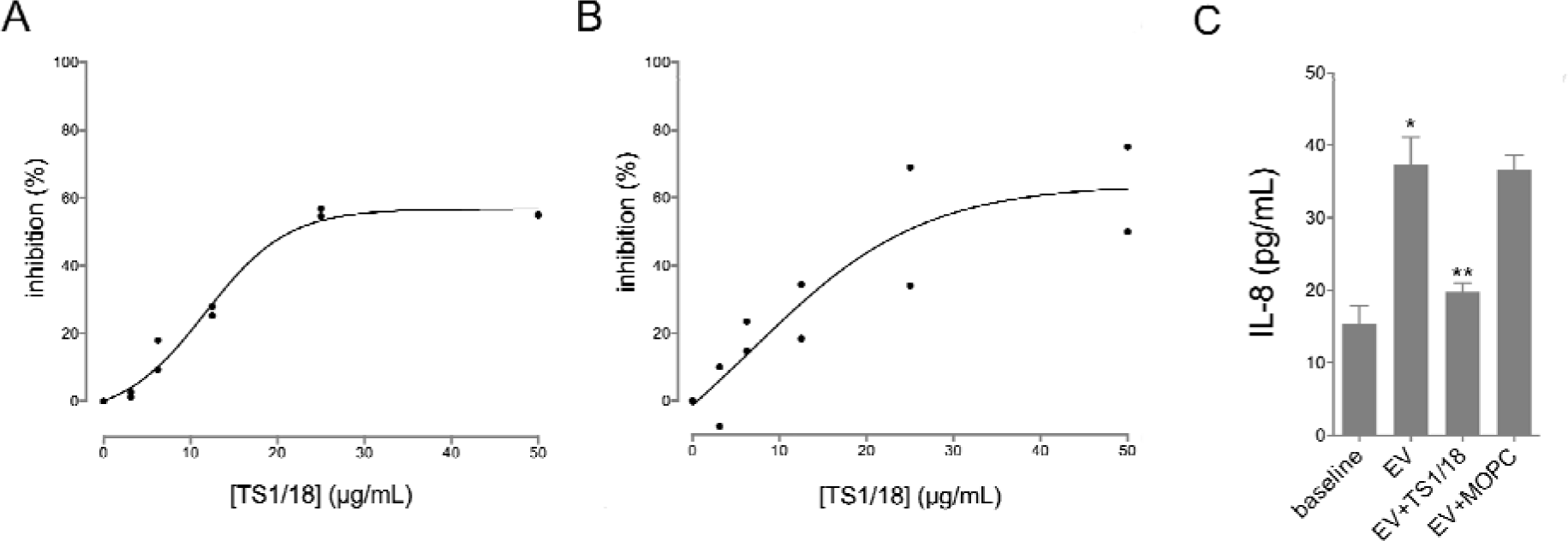
The anti-CD18 monoclonal antibody, TS1/18, inhibits EV-induced synthesis of proinflammatory mediators. EV were incubated with unstimulated lung A549 (A and B) and 16HBE (C) cells for 18 hours, and the conditioned medium tested for the presence of IL-8 (A and C) and MCP-1 (B). * p<.05 for EV-stimulated vs. unstimulated 16HBE cells. ** p<.05 for EV-stimulated 16HBE cells in the absence and in the presence of TS1/18 (25 mg/mL); repeated measures ANOVA. Data from one experiment representative of 8 and 5 for panels A and B, respectively; data form 3 consecutive experiments for panel C.

**Figure 7.**
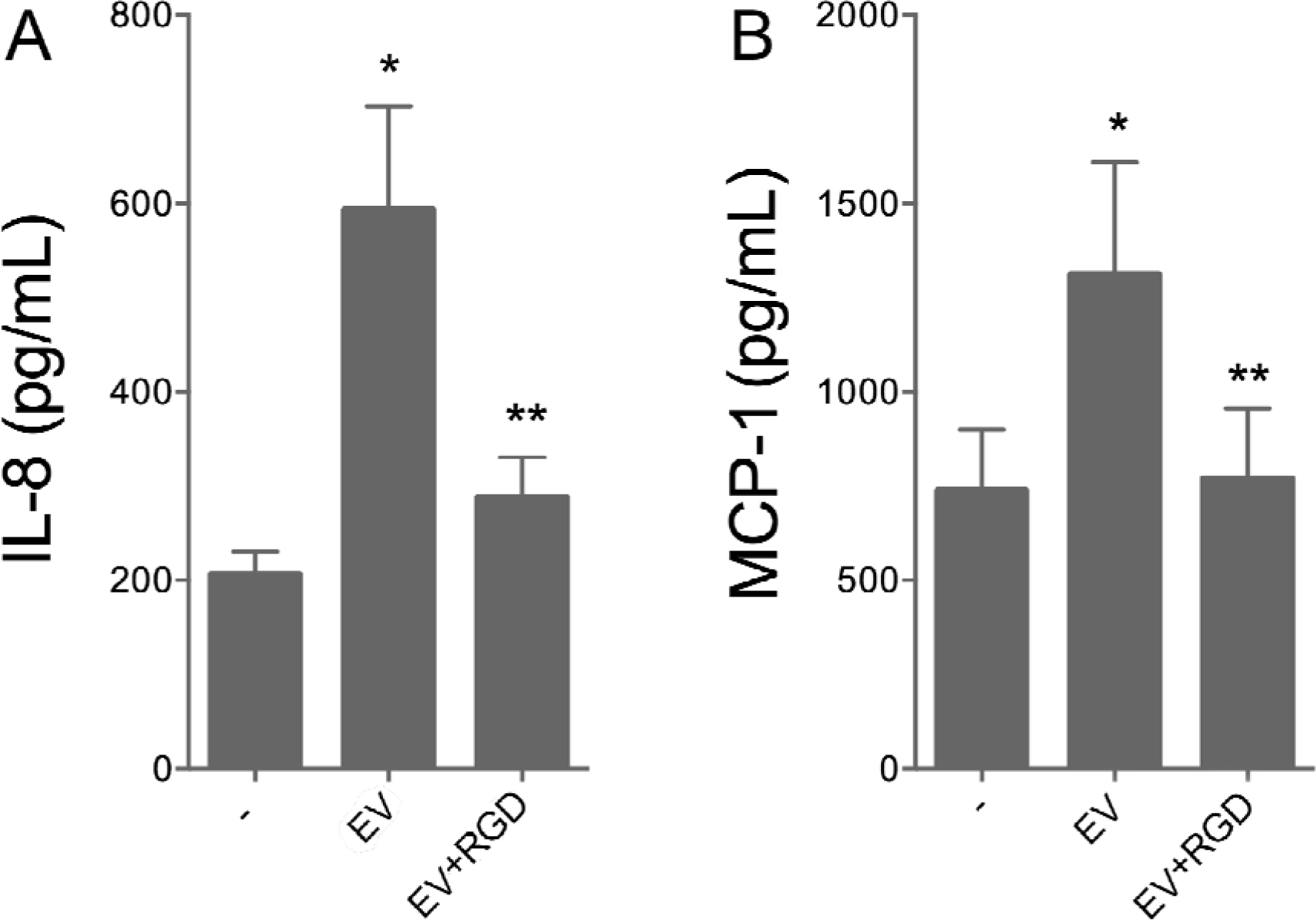
An RGD-containing peptide inhibits EV-induced synthesis of proinflammatory mediators. EV were incubated with unstimulated lung A549 cells for 18 hours, and the conditioned medium tested for the presence of IL-8 (A) and MCP-1 (B). *: p<.05 for EV-stimulated vs. unstimulated A549 cells. ** p<.05 for EV-stimulated A549 cells in the absence and in the presence of the RGD peptide (10 nM); repeated measures ANOVA. Data from 6 and 4 consecutive experiments for panels A and B, respectively.

### EV induce TNF-α expression by A549 cells

Our previous data showing the proinflammatory effects of EV (Cerri et al., 2006) were obtained in experiments performed with epithelial cells not prestimulated with TNF-α. However, the experiments were carried out at late time points (18 h), when the current data show that EV adhere even to unstimulated cells. We hypothesized that an initial, low-level binding of EV upregulates TNF-α synthesis, so that at late time points adhesion takes place even in the absence of external stimuli. Fig. 8 shows that, indeed, TNF-α expression is increased upon incubation of A549 cells with EV.

**Figure 8.**
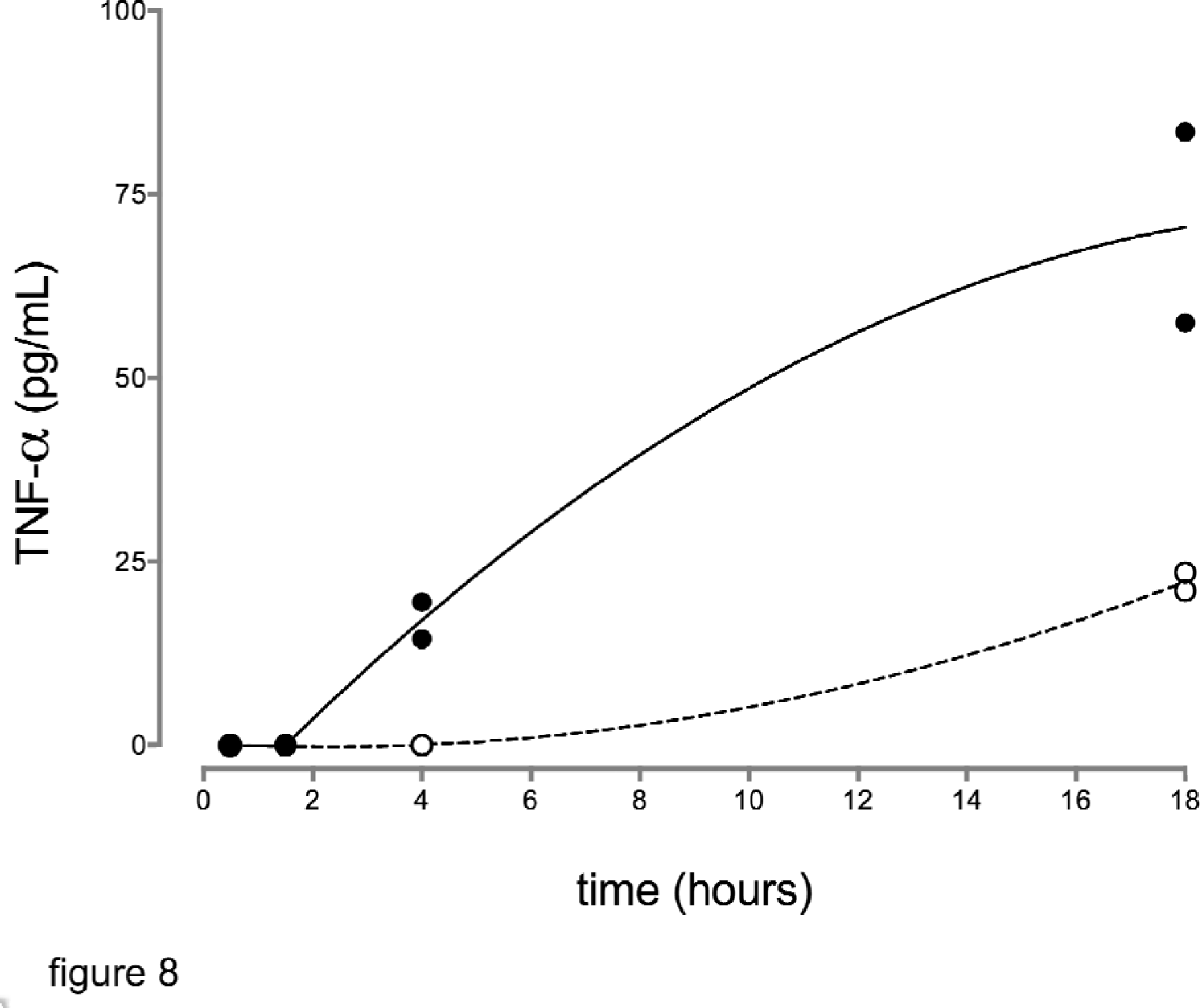
Time-course of EV-induced upregulation of TNF-α synthesis by lung epithelial cells. A549 cells were cultured in the presence (solid line) or absence (dashed line) of EV. Data from one experiment representative of 2

## Discussion

The first aim of this study was to investigate the hypothesis that mononuclear cell-derived EV adhere to bronchial and alveolar epithelia, thereby mimicking the behaviour of their parental cells. Our data indicate that fluorescently labelled EV adhere minimally to lung epithelial cells in baseline conditions and that adhesion is significantly upregulated by stimulation of the latter with the classical proinflammatory agonist, TNF-α. The next step was to investigate which adhesion receptor(s) potentially expressed on the surface of EV is (are) involved in such phenomenon. CD18 is the common β2 chain of the leukocyte integrins, CD11a/CD18 (also known as leukocyte function-associated antigen 1, LFA1), CD11b/CD18 (Mac1), and CD11c/CD18 (p150,95). The role of CD18-mediated adhesion of leukocytes to lung epithelial cells is well established (Celi et al., 1999; Rosseau et al., 2000). Our data, showing that blocking CD18 with a monoclonal antibody prevents EV adhesion, are consistent with the hypothesis that this molecule represents the adhesion receptor involved in such phenomenon. However, since inhibition levels off at approximately 60%, it is possible that other receptors are also involved. CD18 binds to different cognate ligands (Tan, 2012). Of note, metal ion and energy requirement are different for different ligands. Our observation that either Ca^++^ or Mg^++^ is sufficient to support EV adhesion apparently rules out ICAM-1, currently considered one of the most important molecules involved in leukocytes-lung epithelial cells adhesion, as the relevant ligand in this experimental setting (Dransfield et al., 1992); consistent with this hypothesis, EV adhesion was inhibited by a soluble peptide comprising the recognition sequence arginine-glycine-aspartic acid, not present in ICAM-1 (Issekutz et al., 1999). Our group has demonstrated the presence of an ICAM-1-independent, CD18-dependent mechanism of leukocyte adhesion to lung epithelial cells (Celi et al., 1999); however, the metal ions requirements are different from those described in the present data.

Next, we tested the hypothesis that adhesion of EV to lung epithelial cells is required for the proinflammatory effects of the former that has been previously demonstrated. Our data show that when EV adhesion is inhibited either by an anti-CD18 antibody or an RGD - containing peptide, the synthesis of IL-8 and MCP-1 induced by EV is also inhibited, thus lending support to the hypothesis. Again, as inhibition of IL-8 and MCP-1 synthesis parallels that of EV adhesion and levels off at 60%, the presence of other mechanisms, possibly independent of CD18 and/or RGD-containing molecules cannot be ruled out.

The role of adhesion molecules in the recruitment of blood-borne leukocytes to sites of inflammation has generated tremendous interest since the discovery of leukocyte integrins and selectins in the late 1980’s; the model of this so-called “adhesion cascade” has been later refined but remains valid (Ley et al., 2007). The lung epithelium represents an active participant of the innate immune system. As part of the defence mechanisms against pathogens, blood-borne leukocytes may be recruited into the airways through relatively well defined mechanisms involving chemoattractants and adhesion receptors (Khair et al., 1996). Numerous stimuli have been shown to induce the synthesis of such molecules during an inflammatory reaction, including bacterial products (Khair et al., 1996), cigarette smoke, particulate and gaseous pollutants (Chung and Adcock, 2008). EV are increasingly recognized as effectors in relevant biological phenomena. EV can be induced by all the above-mentioned noxious agents (Bernimoulin et al., 2009; Li et al., 2010; Cordazzo et al., 2014; Neri et al., 2016b) and cause the synthesis of molecules involved in the development of an inflammatory reaction. Based on the current data and previous reports, we propose a model whereby mononuclear cell-derived EV, generated upon stimulation with different noxious agents, adhere to the lung epithelium through a CD18-/RGD-mediated mechanism and promote the NFκB-mediated upregulation of the synthesis of proinflammatory agonists. As a dysregulation of such mechanisms is responsible for numerous respiratory diseases (Chung, 2005; Gwyer Findlay and Hussell, 2012), targeting EV generation and adhesion to lung epithelia has the potential to represent a new avenue in the quest for novel therapeutic strategies.

## Materials and Methods

### Reagents

RPMI 1640 medium, MEM, HEPES solution, penicillin/streptomycin, L-glutamine, trypsin/EDTA, calcium ionophore A23187, trypsin, sodium azide, Dulbecco phosphate buffered saline with and without calcium/magnesium, Dextran T500, IgG_1_ isotype control from murine myeloma clone MOPC-21, foetal bovine serum (FBS), the peptides GRGDNP and GRADSP, CaC1_2_, MgCl_2_, and MnCl_2_ were obtained from Sigma (Milan, Italy). LymphoPrep was purchased from Voden Medical Instruments spa (Meda, Italy). Carboxyfluorescein diacetate succinimidyl ester (CSFE) was purchased from Miltenyi Biotec (Calderara di Reno, Italy). Tumor necrosis factor (TNF)-α was purchased from

PeproTech EC Ltd. (London, UK). MA1810 clone number TS1/18 was purchased from ThermoScientific (Rockford, USA). μ-Slide 4 Well were purchased from Ibidi GmbH (Martinsried, Germany). Human TNF-α Elisa Ready-SET-Go!, Human IL-8 Elisa Ready-SET-Go! and Human MCP-1 Elisa Ready-SET-Go! were purchased from Affimetrix eBioscience (San Diego, USA).

### Cell Culture

Cells of the human alveolar epithelial line, A549 (American Type Culture Collection, CCL-195), were kindly provided by Dr. R. Danesi, University of Pisa, Pisa, Italy. A549 cells were maintained in RPMI supplemented with 10% (vol/vol) FBS, 0.2 mg/mL L-glutamine,100 U/mL penicillin, and 100 μg/mL streptomycin. The immortalized bronchial epithelial cells, 16HBE (American Type Culture Collection, CRL-2741) were kindly provided by Dr. M. Profita (National Research Council, Palermo, Italy).16HBE cells were maintained in MEM supplemented with 10% (vol/vol) FBS, 0.2 mg/ml L-glutamine and 2.5 mM HEPS. Both cell lines were maintained in a humidified 95% air/5% CO_2_ atmosphere at 37°C.

### Mononuclear cell isolation and EV generation

Mononuclear cells were isolated either from fresh buffy coats obtained from the local blood bank or from the peripheral blood of normal volunteers as described (Cordazzo et al., 2013) with minor modifications introduced over time in our routine procedures; specifically, Lymphoprep was used instead of Ficoll-Paque; the concentration of dextran was increased to 2.5% and cells were washed with PBS rather than PBS/EDTA. For EV generation, A23187 (12 μM) was added; after 15 min at 37°C, the supernatant was recovered and cleared by two differential centrifugations, the first at 1500 x *g* for 15 min at 4°C to remove dead cells and big cell fragments and the second at 16000 x *g* for 45 min at 4°C to obtain a EV pellet. The pellet was resuspended in PBS and used either in adhesion experiments, or as a stimulus for cytokine expression by epithelial cells. The typical yield was 89 ± 79 (mean ± SD) nM phosphatidylserine, as assessed by a commercially available kit (Zymuphen, Hyphen BioMed, Neuville-sur-Oise, France) as described (Neri et al., 2016a). For experiments designed to assess adhesion to epithelial cells, EV were labelled with CFSE. CFSE is a cell permeant, non-fluorescent pro-dye that is cleaved by intracytoplasmic esterases resulting in an impermeant fluorescent molecule; CFSE labelling therefore allows the differentiation of intact EV from linearized membrane fragments (Grisendi et al., 2015). EV were incubated with CFSE (0.5μM) for 15’, pelleted by centrifugation (16000 x g for 45 min at 4°C) and resuspended in PBS.

### Electron microscopy

Transmission electron microscopy (TEM) was performed on isolated vesicles, resuspended in PBS, to analyze their ultrastructural morphology. Following proper dilutions, the samples were adsorbed to 300 mesh carbon-coated copper grids (Electron Microscopy Sciences, Hatfield, PA, USA) for 5 minutes in a humidified chamber at room temperature. Vesicles on grids were then fixed in 2% glutaraldehyde (Electron Microscopy Sciences, Hatfield, PA, USA) in PBS for 10 minutes and then briefly rinsed in milli-Q water. Grids with adhered vesicles were examined with a Philips CM 100 transmission electron microscope TEM at 80kV, after negative staining with 2% phosphotungstic acid, brought to pH 7.0 with NaOH. Images were captured by a Kodak digital camera.

### EV adhesion assay to lung epithelial cells

A549 and 16HBE were plated on Ibidi μ-Slide 4 Well at 210^5^ cells/well. When confluent, cells were washed once with PBS and incubated with TNF-α (25 ng/ml) or vehicle for 18h. The cells were then washed three times with PBS with Ca^2^+ and Mg^2^+, taking great care to preserve the integrity of the monolayers. EV were incubated with A549 and 16HBE cells for 1,5h and then washed gently three times. Adhesion experiments with A549 cells were carried out in PBS, while 16HBE cells required culture medium to avoid cell detachment. At the end of the assay, the plates were analysed by fluorescence microscopy to enumerate EV using a Nikon Eclipse Ti microscope (Amsterdam, The Netherlands), final magnification 200x.

In experiments designed to evaluate the role of different metal ions, EV were resuspended in PBS containing either Ca^2^+ only or Mg^2^+ only or Mn^2^+, and the cell monolayers were washed throughout the test with PBS containing the same divalent ion.

In experiments designed to understand the requirements for metabolic energy, the adhesion assay was carried out at 4 °C or in the presence of 10 mM NaN_3_.

In experiments designed to understand if EV adhesion to lung epithelial cells is mediated by the integrin β_2_ subunit, CD18, EV were preincubated for 30’ with the inhibitory anti-CD18 monoclonal antibody, TS1/18.

### Cytokine assays

The conditioned medium of epithelial cells incubated with EV under the experimental conditions described was analysed for IL-8, MCP-1 and TNF-α content with commercially available ELISA kits following the manufacturers’ directions.

### Data presentation and statistical analysis

Data are shown as mean±SEM; comparisons among groups were made by either ANOVA for repeated measurements followed by Bonferroni’s analysis or paired t-test as appropriate; data were analyzed using Prism Software (GraphPad, San Diego, CA, USA). All tests were two-tailed. Values of p<0.05 were considered statistically significant.

## Competing interests

The Authors declare no competing interests.

## Fundings

This research received no specific grant from any funding agency in the public, commercial or not-for-profit sectors.

